# Cortisol levels after cold exposure are independent of adrenocorticotropic hormone stimulation

**DOI:** 10.1101/670836

**Authors:** Alissa Shida, Tomoya Ikeda, Naoto Tani, Fumiya Morioka, Yayoi Aoki, Kei Ikeda, Miho Watanabe, Takaki Ishikawa

## Abstract

We previously showed that postmortem serum levels of adrenocorticotropic hormone (ACTH) were significantly higher in cases of hypothermia (cold exposure) than other causes of death. This study examined how the human hypothalamic-pituitary-adrenal axis, and specifically cortisol, responds to hypothermia.

**Human samples:** Autopsies on 205 subjects (147 men and 58 women; age 15-98 years, median 60 years) were performed within 3 days of death. Cause of death was classified as either hypothermia (cold exposure, *n*=14) or non-cold exposure (controls; *n*=191). Cortisol levels were determined in blood samples obtained from the left and right cardiac chambers and common iliac veins using a chemiluminescent enzyme immunoassay. Adrenal gland tissue samples were stained for cortisol using a rabbit anti-human polyclonal antibi.

**Cell culture:** AtT20, a mouse ACTH secretory cell line, and Y-1, a corticosterone secretory cell line derived from a mouse adrenal tumor, were analyzed in mono-and co-culture, and times courses of ACTH (in AtT20) and corticosterone (in Y-1) secretion were assessed after low temperature exposure mimicking hypothermia and compared with data for samples collected postmortem for other causes of death. However, no correlation between ACTH concentration and cortisol levels was observed in hypothermia cases. Immunohistologic analyses of samples from hypothermia cases showed that cortisol staining was localized primarily to the nucleus rather than the cytoplasm of cells in the zona fasciculata of the adrenal gland. During both mono-culture and co-culture, AtT20 cells secreted high levels of ACTH after 10-15 minutes of cold exposure, whereas corticosterone secretion by Y-1 cells increased slowly during the first 15-20 minutes of cold exposure. Similar to autopsy results, no correlation was detected between ACTH levels and corticosterone secretion, either in mono-culture or co-culture experiments. These results suggested that ACTH-independent cortisol secretion may function as a stress response during cold exposure.

## Introduction

Many reports have documented the pathologic changes observed in humans affected by hypothermia due to cold exposure, and “classic” morphologic findings supporting a diagnosis of hypothermia have been established [1-7]. However, as other etiologies of hypothermia include drug abuse, dementia, malnutrition, and infectious disease, only a few studies have specifically examined pathologic findings after cold exposure [8,9], especially from a biochemical perspective, such as the presence and levels of ketone bodies [10-13]. Furthermore, only a few reports have estimated hormone levels as part of the pathophysiologic findings of cold exposure [14-16].

The primary stress response system is the sympathetic/adrenomedullary (S/A) system, which includes the chromogranin A [14] and hypothalamic-pituitary-adrenal (HPA) axis [16, 18]. Previous studies have suggested that postmortem serum adrenocorticotropic hormone (ACTH) concentration is a useful biomarker of death due to cold exposure and the magnitude of physical stress responses during cold exposure [17]. Increased serum concentrations of ACTH associated with activation of the HPA axis and S/A system can be biochemically evaluated by measuring catecholamine and chromogranin A levels [19-23]. With respect to the HPA axis, it is known that cortisol levels are correlated with ACTH levels, and a precursor of cortisol, which is an activator, also inactivates cortisone accounting for 4-5% and corticosterone exhibiting only weak activity [24, 25]. Thus, this study evaluated cortisol as a biomarker of cold exposure-related stress by analyzing cases of human death due to hypothermia. We also assessed the relationship between ACTH and corticosterone levels during cold exposure using a mouse cell culture model.

## Material and Methods

### Autopsy samples

Autopsies were performed within 3 days postmortem at our institute. The study included 205 serial cases (147 men and 58 women), and the median age was 60 years (range 15-98 years). Cortisol levels were determined in blood samples collected aseptically from the left and right cardiac chambers and the common iliac vein using syringes.

Cause of death was determined based on findings from a complete autopsy as well as macromorphological, micropathologic, and toxicologic examinations. Cases were classified as either hypothermia (cold exposure, *n*=14) or control. Cause of death in the latter group included blunt injury (*n*=37 total; head injury [*n*=28], non-head injury [*n*-=9]), sharp-instrument injury (*n*=8), fire fatality (*n*=43), asphyxia (*n*=28), intoxication (*n*=12 total; methamphetamine-related fatality [*n*=3], psychotropic drugs [*n*=6], other [*n*-=3]), drowning (n=12), hyperthermia (heat stroke, *n*=10), acute ischemic heart disease (*n*=20), and natural causes (*n*=22). Case profiles are shown in Table1.

**Table1.** Case profile.

| Cause of Death | Number | Gender (male/female) | Age-structure (mean) | Survival period (mean) | Postmortem period (mean) | Hospitalization (male/female) |
| --- | --- | --- | --- | --- | --- | --- |

|  |  |  |  | n,<br>h)α | h)α |  |
| --- | --- | --- | --- | --- | --- | --- |
| Hypothermiaα | 14α | 9/5α | 34-89¶<br>(62)α | 6-24¶<br>(18)α | 24-72¶<br>(52.6)α | 0/0α |
| Blunt injury ¶<br>(head injury)α | 28α | 19/9α | 15-98¶<br>(66)α | <0.5-1056<br>¶<br>(128.<br>7)α | 12-60¶<br>(29.3)α | 9/6α |
| Blunt injury¶<br>(non-head injury)α | 9α | 9/0α | 52-85¶<br>(67)α | <0.5-960¶<br>(122.<br>6)α | 24-60¶<br>(30.6)α | 4/0α |
| Sharp instrument injuryα | 8α | 7/1α | 40-85¶<br>(67)α | <0.5-24¶<br>(6.3)<br>α | 12-36¶<br>(27.4)α | 3/1α |
| Fire fatalityα | 43α | 34/10α | 28-95¶<br>(73)α | <0.5-3600<br>¶<br>(142.<br>4)α | 12-60¶<br>(27.8)α | 6/2α |
| Asphyxiaα | 29α | 19/10α | 21-83¶ | <0.5- | 12-60¶ | 4/2α |
|  |  |  | (57)☐ | 240☐<br>(23.7)<br>☐ | (33.4)☐ |  |
| Intoxication<sup>a</sup>☐ | 11☐ | 8/3☐ | 25-59☐<br>(38)☐ | <0.5-48☐<br>(11)☐ | 12-36☐<br>(32.7)☐ | 0/1☐ |
| Drowning☐ | 11☐ | 7/4☐ | 44-85☐<br>(62)☐ | <0.5-2☐<br>(3)☐ | 12-48☐<br>(29.6)☐ | 0/0☐ |
| Hyperthermia☐ | 10☐ | 3/7☐ | 28-92☐<br>(70)☐ | 6-240☐<br>(33.1)<br>☐ | 24-48☐<br>(32.7)☐ | 2/1☐ |
| Acute ischemic heart disease☐ | 20☐ | 19/1☐ | 19-88☐<br>(61)☐ | <0.5-144☐<br>(16.5)<br>☐ | 6-60☐<br>(33.6)☐ | 1/1☐ |
| Other natural death☐ | 22☐ | 14/8☐ | 21-88☐<br>(70)☐ | <0.5-4320<br>☐<br>(243.<br>5)☐ | 24-48☐<br>(29.2)☐ | 5/3☐ |

<sup>a</sup>Methamphetamine-related fatalities, n=3; psychotropic drugs, n=5; others, n=3

Cases of hypo- and hyperthermia due to drug abuse and bathing, respectively, were excluded. Postmortem interval was defined as time elapsed from estimated time of death to autopsy, whereas survival period was defined as the time from the onset of fatal insult to death. Only clearly described cases were examined in this study.

Tissue specimens of the bilateral adrenal glands were collected and fixed in 4% paraformaldehyde in phosphate-buffered saline (PBS; pH 7.2) for histopathologic and immunohistochemical analyses.

### Biochemical analysis

Blood samples were immediately centrifuged to prepare serum, and ACTH and cortisol levels were measured using an AIA-360^®^analyzer (TOSOH Bioscience GmbH, Griesheim, Germany) [27,28]. This analyzer utilizes a competitive fluorescent enzyme immunoassay format and is performed entirely within small, single-use test cups containing all necessary reagents. The analyte in the sample competes with the enzyme-labeled hormone and incubated with a fluorogenic substrate, 4-methylumbelliferyl phosphate. The amount of enzyme-labeled hormone that binds to the beads is inversely proportional to the hormone concentration in the test sample. Calibration, daily checks, and maintenance procedures were carried out as described in the Systems Operator’s Manual. Accurate performance data for human ACTH and cortisol, including analyte recovery and dilution studies, had been previously evaluated and were available in the manufacturer’s technical bulletins. The time required to obtain the first result using this assay is 20 minutes, with additional results obtained every minute thereafter.

Serum samples (150μL each) were placed in the test cups, and both hormones were measured using the above-mentioned immunoassays. The lower (and upper) reported values for the ACTH and cortisol assays were 2.0 (2000.0) pg/mL and 28.0 (1656.0) nmol/L, respectively.

### Oxyhemoglobin measurement

Blood oxyhemoglobin was determined using a CO-oximeter system (ABL80FLEX System; Radiometer Corp., Tokyo, Japan) in hypothermia patients [29, 30]. Blood alcohol levels were determined using headspace gas chromatography/mass spectrometry (GC/MS), and amphetamine and psychotropic drugs were detected by GC/MS [17].

### Immunohistochemistry

Harvested adrenal glands were fixed in 4% paraformaldehyde in PBS (pH 7.2) for 12 h, embedded in paraffin, and sectioned at a thickness of 4μm. Deparaffinization (Sakura tissue TEK DRS 2000, Tokyo, Japan) of each section was followed by heat-mediated antigen retrieval in citrate buffer (pH7.0) for 10min, after which each section was immersed in 0.3% H_2_O_2_-methanol for 10 min to inactivate endogenous peroxidases. After washing in PBS for 5 min, slides were incubated overnight with anti-cortisol-binding globulin antibody (ab107368; Abcam). Immunoreactivity was visualized by the polymer method using Dako Envision+ Dual Link System-HRP (K4063; Dako, CA, USA) and the Dako liquid DAB+ Substrate Chromogen System (K3468; Dako), according to the manufacturer’s instructions and with hematoxylin counterstaining [13, 17]. The total number of cells in the adrenal gland and number of cells exhibiting cytoplasmic or nuclear cortisol immunoreactivity were determined microscopically under 400× magnification. Three random fields were independently enumerated, and the data are presented as number of cortisol-positive cells (cytoplasm or nucleus, respectively)/total number of adrenal gland cells×100. As cells in the zona fasciculata of the adrenal gland are known to produce cortisol in the cytoplasm, immunostaining for cortisol in each group was evaluated by technicians blinded to sample grouping. Three sections were randomly selected for cell counting [31, 32].

### Cell culture models

#### Mono-culture models of pituitary and adrenal cells

Mono-culture models of ACTH-secreting AtT20 pituitary cells [33-37] and corticosterone-secreting Y-1 adrenal cells [38-42] derived from mice were developed to verify whether these cells secrete hormones only upon stimulation by exposure to cold. For both cell types (AtT20 and Y-1), the culture medium consisted of a 1:1 ratio of DMEM-F12 and 15% charcoal stripped fetal bovine serum (FBS; Biological Industries,CT.,USA) with 4mM L-glutamine, 50 U/mL penicillin, and 50 μg/ml streptomycin. Initially, cells of both types were seeded and cultured at 37°C. Growth was controlled at 54,618 cells/cm^2^ for AtT20 and 57,803 cells/cm^2^ for Y-1, and the cells were allowed to proliferate until they covered the surface of the culture dishes. The culture medium for Y-1 cells was replaced once every 2 days. Once the AtT20 and Y-1 cells reached confluence, they were transferred to 4°C and maintained. The amount of ACTH and corticosterone in the culture medium was measured at 5, 10, 15, 20, 30, 40, 60, 180, and 360 min; at 12 and 24 h; and at 3 and 5 days. ACTH was measured using a mouse ACTH assay kit (FEK-001-21; Phoenix Pharmaceuticals, Inc., USA) [43,44], and corticosterone was measured using a mouse corticosterone assay kit (Assay MAX EC3001-1; ANG, USA) [45-47]. At the end of the experiment, adherent cells were dissociated from the surface using trypsin and then counted; hormone concentrations were calculated using a correction formula and the measured values.

#### Co-culture model development

We developed a co-culture system for AtT20 ACTH-secreting cells (ECACC no. 87021902) [36,37] and Y-1 corticosterone-secreting cells derived from mice [45-47] as a model of the pituitary-adrenal system. The co-culture model was used to investigate whether these cells interact as part of the HPA axis during cold-stimulated hormone production. Both AtT20 and Y-1 cells were cultured in medium containing DMEM-F12 supplemented with 15% inactivated FBS, 50 μg/ml streptomycin, 50 μM penicillin, and 0.25 μg/ml fungizone. To inactivate ACTH included in the culture medium, 0.2 mL of rabbit anti-mouse ACTH (1-24) serum (Siemens, Immulyze) was added to 200 mL of culture medium, we decided about the proper amount of the rabbit serum using an ACTH ELISA kit (MDB, M046006) [48].

Initially, both AtT20 and Y-1 cells were cultured separately at 37°C, with AtT20 and Y-1 cells on the top and bottom of the filter, respectively. The cells were then co-cultured at 4°C. The insert for 6-well plate (Greiner Bio-One, Frickenhausen, Germany) used to separate the AtT20 and Y-1 cells had a diameter of 23.1 mm, pore size of 3.0 μm, and pore density of 2×10^6^ pores/cm^2^. Initially, corticosterone-secreting Y-1 cells were cultured on the bottom of the filter with the filter placed upside down so that the cells formed a mono layer. Subsequently, the filter was placed upright in the culture medium. Schroten H, (2016) established this method in a choroid plexus model [49-53], and we previously described this method in a report on the physiologic significance of the blood-cerebrospinal fluid barrier and prolactin [54].

Excessive growth on the filter was controlled by trypsinization to maintain a single layer of cells; the number of Y-1 cells on the filter was limited to 57,803/cm^2^. As the Y-1 cells formed tights junctions, movement of ACTH between the cells was prevented. Thereafter, the culture medium was replaced once every 2 days. Once culturing of the Y-1 cells was complete, AtT20 (ACTH secreting) cells were similarly grown on the other side of the filter (i.e., the side opposite to Y-1 cells). The filter was immersed in the culture medium by placing the ACTH-secreting (AtT20) cells side facing up and corticosterone-secreting (Y-1) cells side facing down. Levels of ACTH and corticosterone in the culture medium were measured at 5, 10, 15, 20, 30, 40, and 60 min, as indicated above. After measurement of both hormones, the adherent cells were dissociated from the filter using trypsin and counted. Accurate hormone concentrations were calculated using a correction formula and the measured values.

### Statistical analysis

For comparisons between groups, we used the nonparametric Mann-Whitney *U* test. The Games-Howell test was used for analyses involving multiple comparisons. All analyses were performed using Microsoft Excel and IBM SPSS statistic viewer 24. Lines in each box represent the median, whereas lines outside each box represent the 90% confidence interval. The sensitivity and specificity for distinguishing between two groups using cut-off cortisol values based on blood collection site (i.e., left and right cardiac chambers and common iliac veins) were estimated using receiver operating characteristic (ROC) curve analysis. Areas under the curve were calculated and analyzed using a 1-tailed test. The optimal compromise between sensitivity and specificity was determined graphically.

### Ethics statement

This study was evaluated by the Independent Ethics Committee of the Osaka City University Graduate School of Medicine, which approved opt-out for informed consent regarding the autopsy data analysis (authorization no.4153).

## Results

### Relationship between cortisol levels and sex, age, survival period, and postmortem period

Serum cortisol levels were not associated with postmortem period, survival period, sex and related differences, or age.

### Relationship between cortisol levels and collection site

Cortisol levels exhibited correlation (*R*=0.63-0.92) with blood collection site, namely, left and right cardiac chambers and external iliac vein.

### Relationship between cortisol levels and cause of death

At all blood collection sites, cortisol levels were approximately three times higher in hypothermia cases than in cases involving other causes of death (*p*<0.05-*p*<0.0001; Fig 1a-c). Specifically, serum cortisol levels were significantly higher in hypothermia cases compared with other causes of death: left cardiac blood, 20-120 μg/dL (median 50 μg/dL); right cardiac blood, 20-100 μg/dL (median 50 μg/dL); iliac vein, 20-130 μg/dL (median 60 μg/dL) versus left cardiac blood, 0-50 μg/dL (median 20 μg/dL); right cardiac blood, 0-40 μg/dL (median 10 μg/dL); iliac vein, 0-20 μg/dL (median 20 μg/dL). Furthermore, most cases exhibited lower cortisol levels, except in hyperthermia cases (heat stroke: left cardiac blood, 0-60 μg/dL [median 30 μg/dL]; right cardiac blood, 0-42 μg/dL [median 20 μg/dL]; iliac vein, 0-60 μg/dL [median 20 μg/dL]). There was no correlation between ACTH concentration and cortisol in hypothermia cases at any of the collection sites tested (left cardiac blood: Y = 0.0103x + 3.237; *r*=0.065; *p*> 0.05 versus right cardiac blood: Y = 0.0217x + 2.7124; *r*=0.113; *p*>0.05 versus iliac vein: Y = 0.026x + 2.4458; *r*=0.170; *p*>0.05).

**Fig 1.**
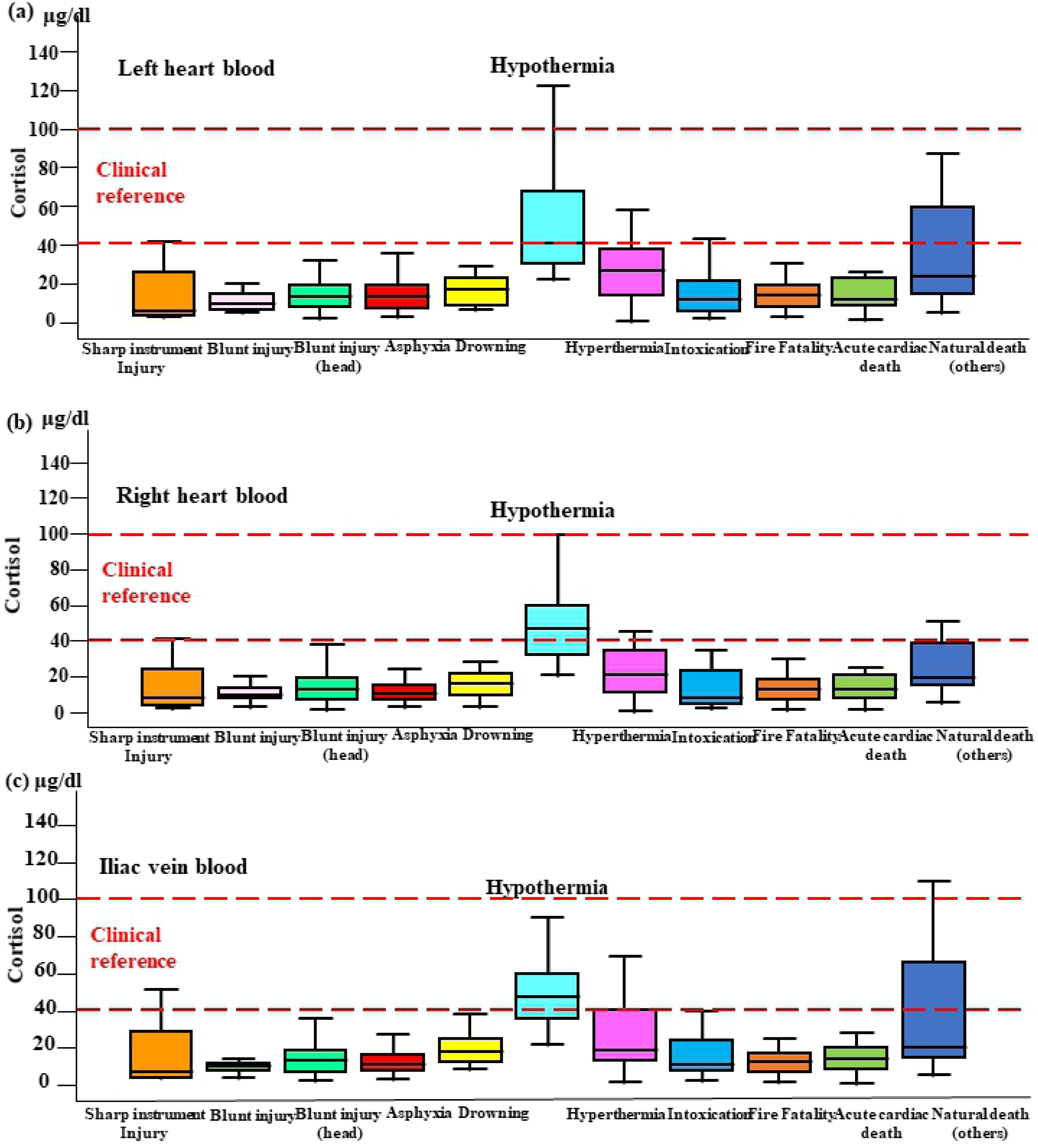
Cortisol levels in blood collected from three sites. Cortisol levels by cause of death in the left (a) and right (b) cardiac chambers and the common iliac vein (c).

Sensitivity and specificity cut-off values for distinguishing between groups with higher (hypothermia) and lower (other cause of deaths) cortisol levels were determined using ROC curve analysis and estimated as 30 μg/mL (0.917 and 0.852) for the left cardiac chamber, 25 μg/mL (0.917 and 0.836) for the right cardiac chamber, and 30 μg/mL (0.917 and 0.872) for the common iliac veins.

### Cortisol immunopositivity in the adrenal gland

Cortisol immunostaining analysis indicated that in hypothermia cases, cortisol was primarily localized in the nucleus, whereas cortisol staining was predominant in the cytoplasm in cases involving other causes of death (Fig 2 a-c). the Graph in Fig 3a shows the cortisol positivity rate in the nucleus by cause of death. Hypothermia (0-70%, median 50%) cases exhibited significantly higher cortisol positivity rate than the other groups (0-30%, median 5%). The graph in Fig 3b shows the number of cells that were positive for cortisol in the cytoplasm; however, it was not significantly different compared with the nucleus.

**Fig 2.**
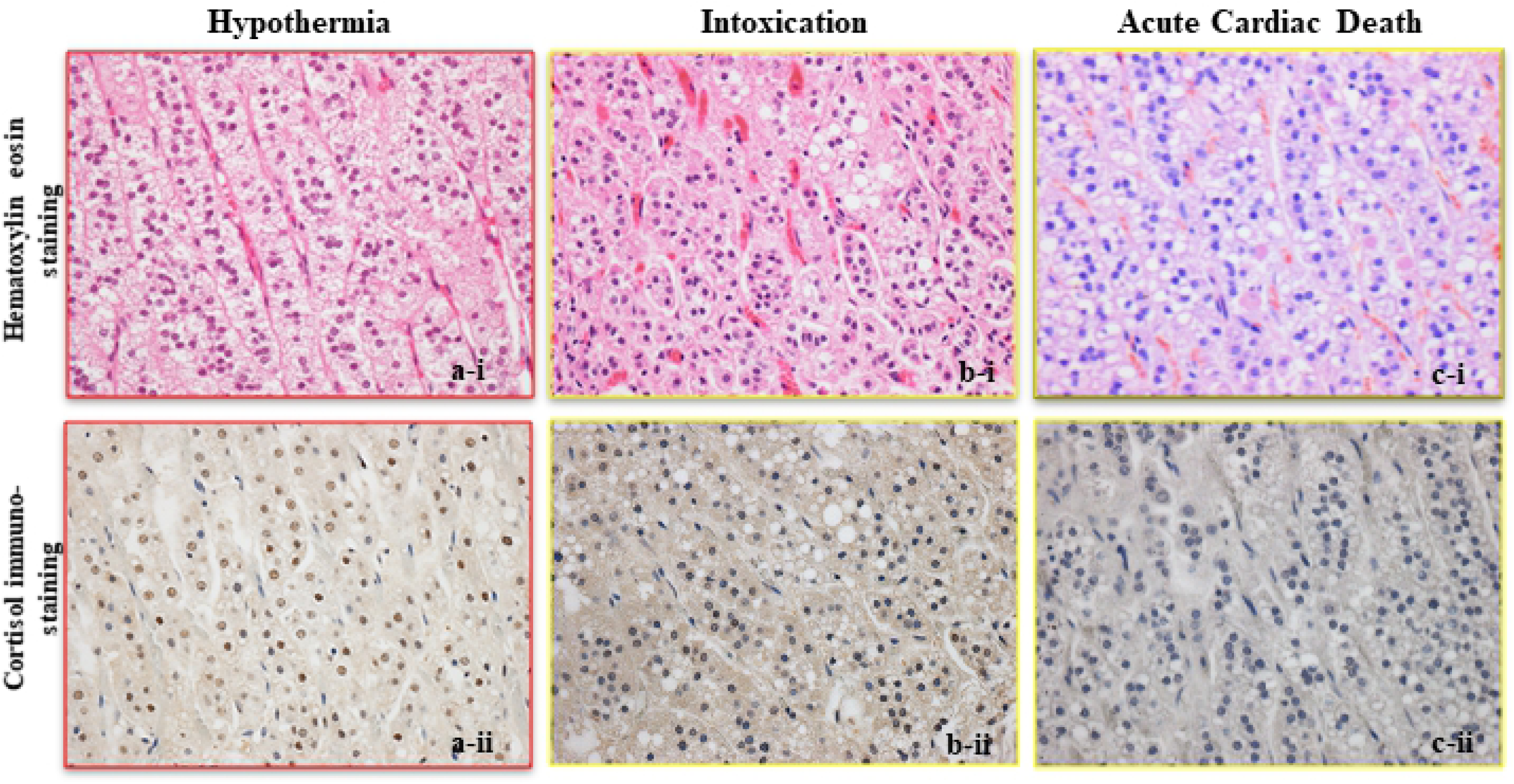
Immunostaining of cortisol in the adrenal gland. Micrographs showing hematoxylin-eosin staining (i) and immunostaining (ii) of cortisol in the adrenal gland in cases of (a) hypothermia, (b) intoxication, and (c) acute cardiac death (original magnification ×100).

**Fig 3.**
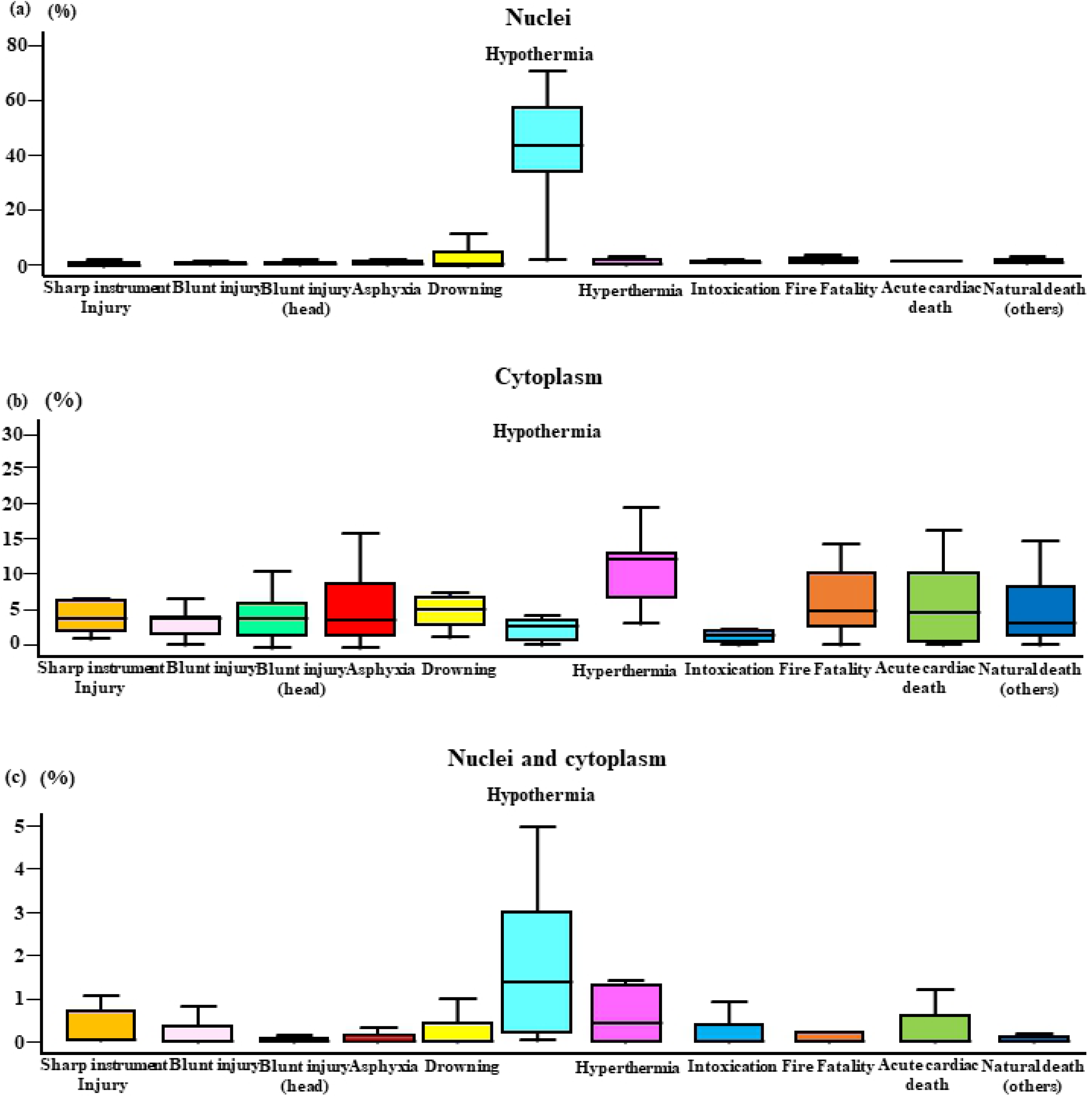
Cortisol positivity rate in the nucleus and cytoplasm by cause of death. Cortisol immunopositivity in the nucleus (a: hypothermia; *p* < 0.05), cytoplasm (b: hypothermia; *p* > 0.05), and nucleus to cytoplasm (c: hypothermia; *p* > 0.05) ratio by cause of death.

### Mono-culture model

In the mono-culture models, ACTH- and corticosterone-secreting cells were cultured separately at 4°C to ensure the absence of ACTH in the culture of corticosterone-secreting Y-1 cells (Fig 4a). AtT20 cells secreted ACTH after 10∼15 min of cold exposure (10 min: median 120 pg/mL; 15 minutes: median 100 pg/mL), which subsequently decreased by 30 min (median 15 pg/mL) (Fig 5a). Corticosterone secretion by Y-1 cells increased slowly during the first 30 min of cold exposure (median 30 ng/mL) and subsequently decreased by 60-180 min (60 min: median 25 ng/mL; 180 min: median 20 ng/mL) (Fig 5b). However, cell culture studies did not reveal a correlation between ACTH and corticosterone secretion in mono-culture experiments, and these results thus suggest that corticosterone secretion after cold exposure is independent of ACTH (Fig 5c).

**Fig 4.**
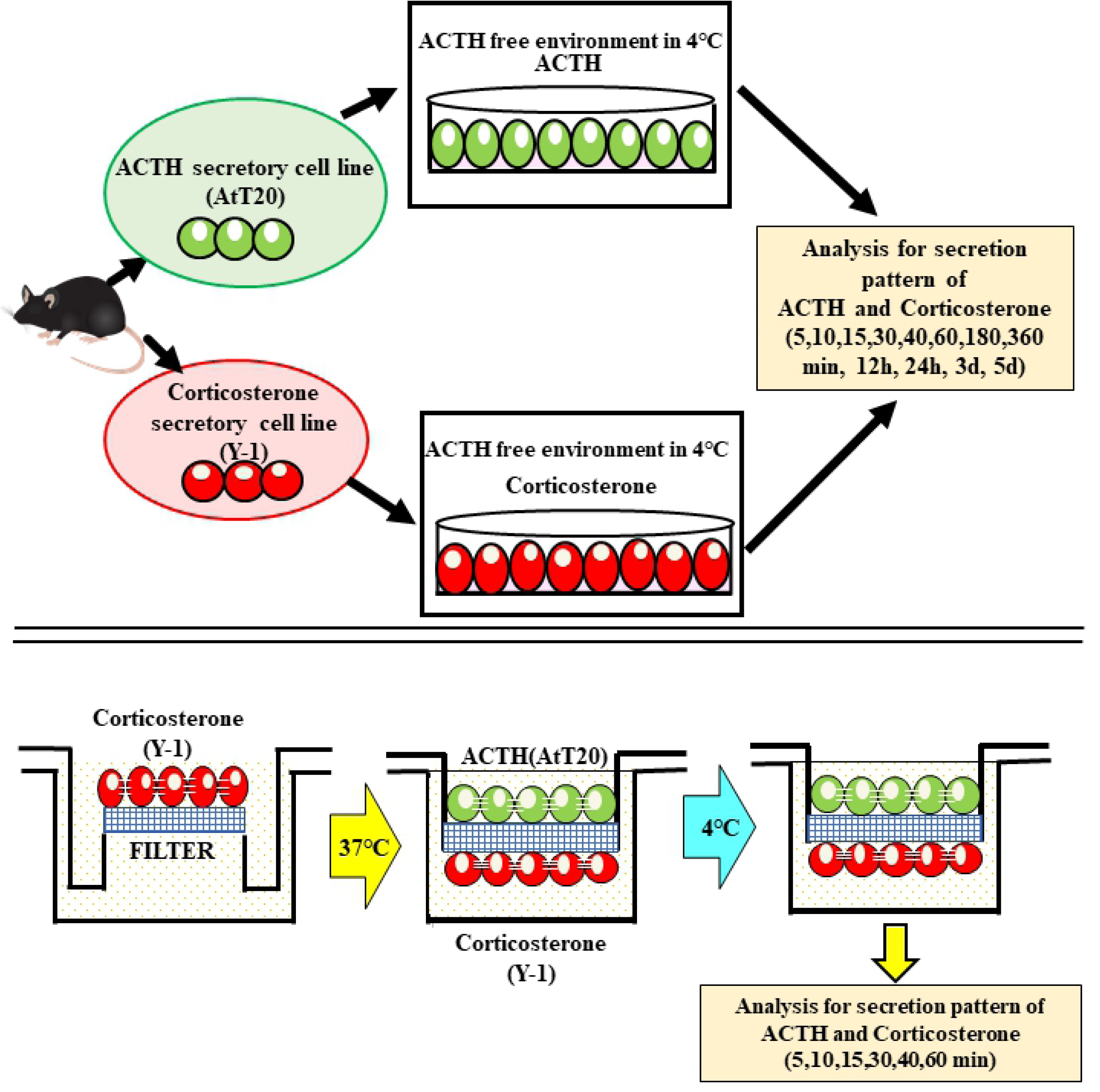
Mono- and co-culture of ACTH- (AtT20) and corticosterone-secreting (Y-1) cells. Schematic illustration of mono-culture (a) and co-culture (b) models of pituitary and adrenal gland cells.

**Fig 5.**
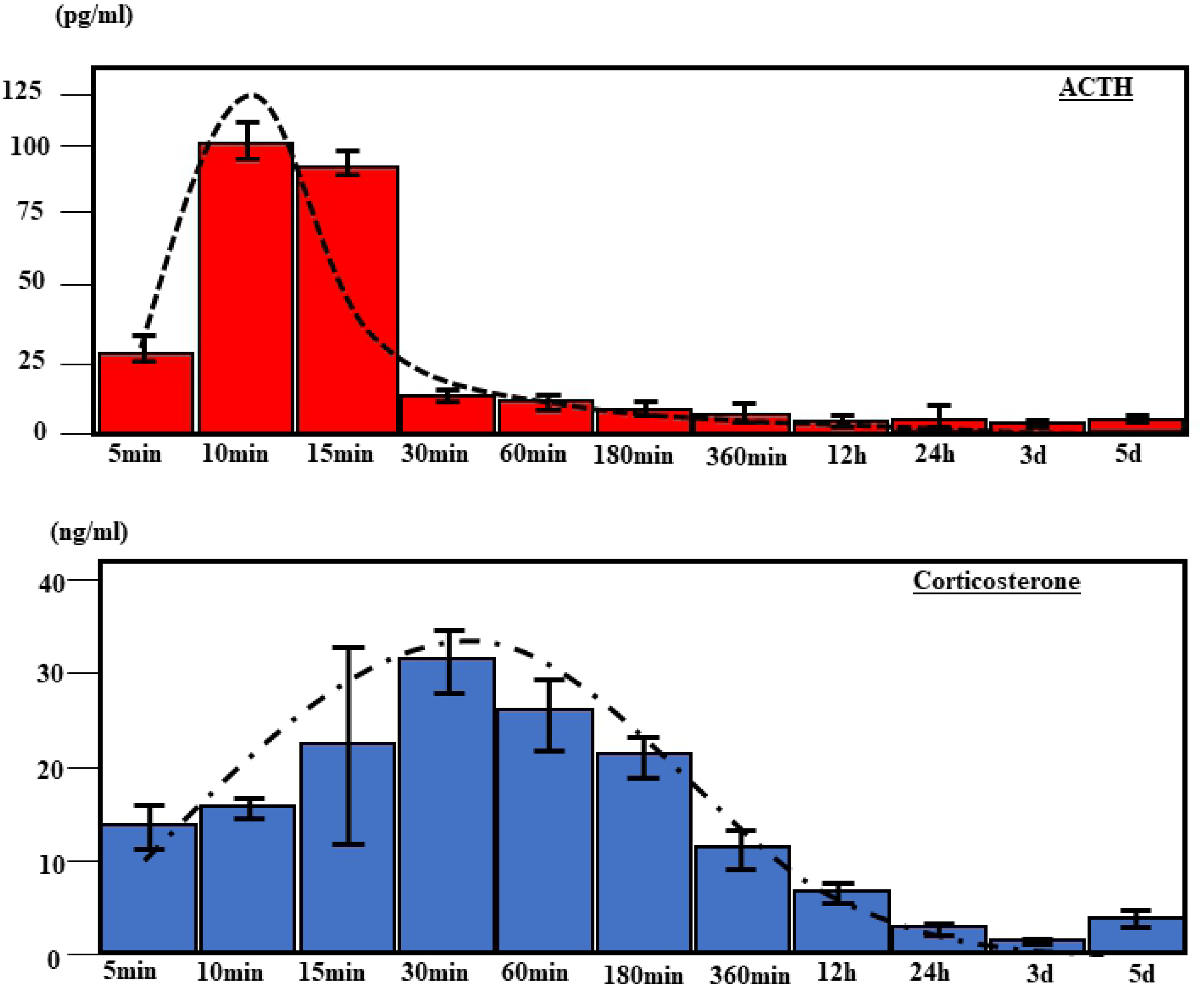
Patterns of ACTH (AtT20) and corticosterone (Y-1) secretion over time. ACTH (a) and corticosterone (b) concentrations over time under cold conditions (4°C) in mono-culture.

### Pituitary–adrenal cell co-culture model

In the co-culture model (Fig 4b), ACTH secretion peaked at 10∼15 min (10 min: median 130 pg/mL; 15 min: median 120 pg/mL) and slowly decreased from 20 min onwards (median 20 pg/mL). Corticosterone levels slowly increased beginning at 10 min (median 30 ng/mL), peaked at 20 min (median 300 ng/mL), and decreased after 30 min (median 150 ng/mL) (Fig 6). These co-culture results suggest that corticosterone secretion is ACTH independent, as seen in mono-culture experiments (Fig 7).

**Fig 6.**
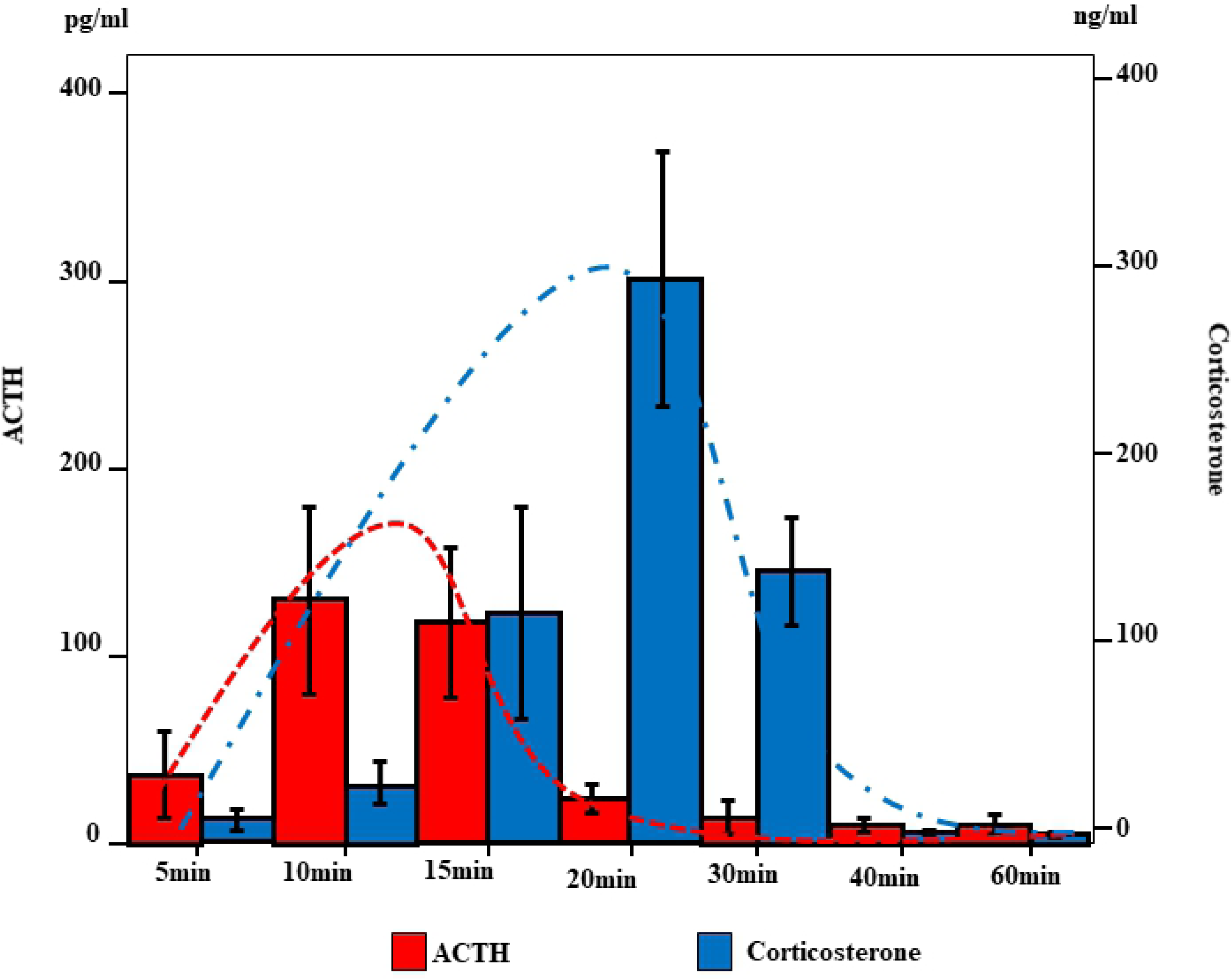
Secretion of ACTH and corticosterone over time in co-culture of AtT20 and Y-1 cells. Concentrations of ACTH and corticosterone over time in co-culture of ACTH-(AtT20) and corticosterone-secreting (Y-1) cells under cold conditions (4°C).

**Fig 7.**
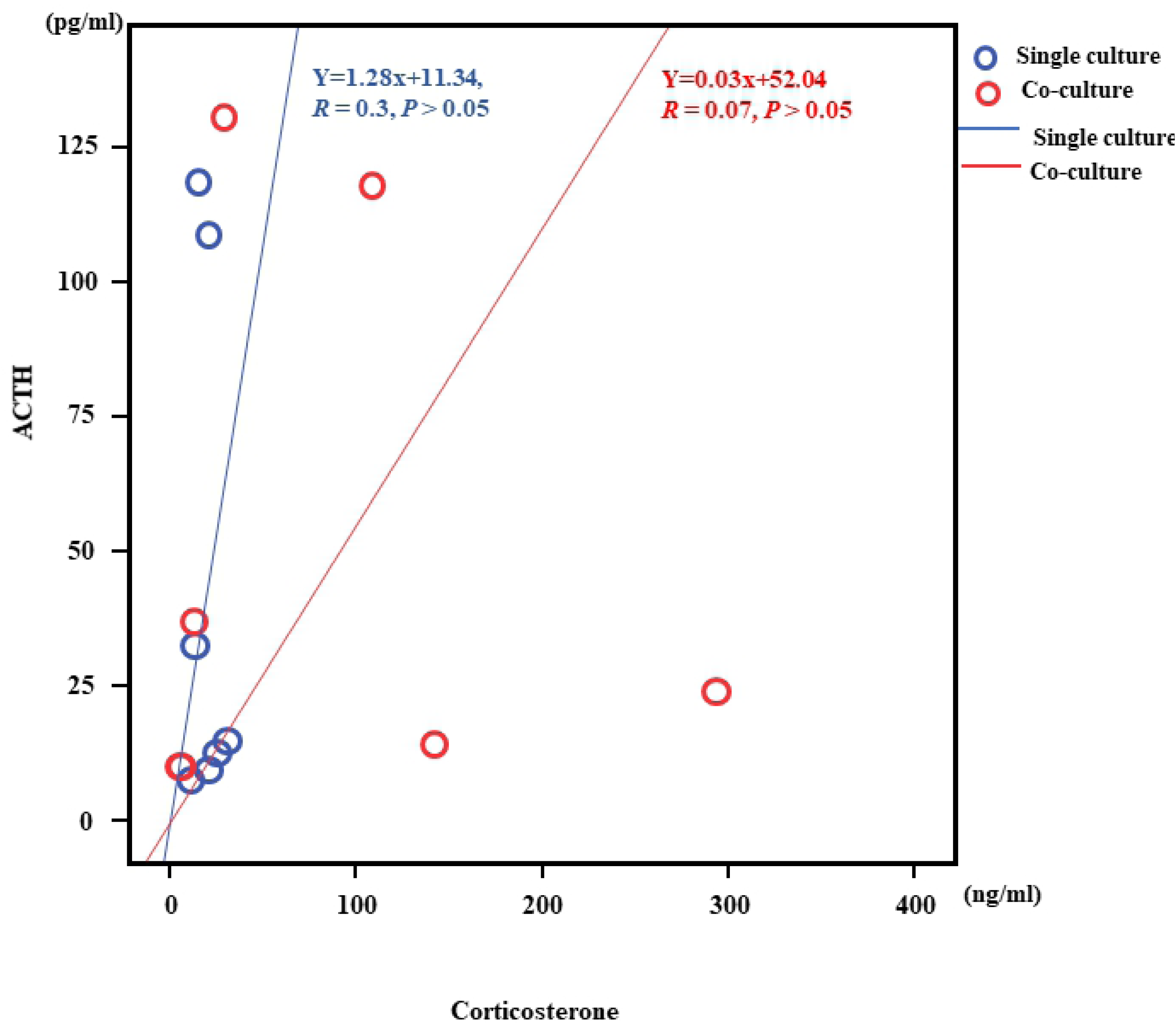
Correlation of ACTH and Corticosterone in mono-culture and co-culture. The correlation between ACTH and corticosterone levels in mono-culture. The mono-culture study demonstrated that corticosterone secretion following cold exposure is independent of ACTH (Y = 1.28x + 11.34, *r* = 0.3, *p* > 0.05). In co-culture the correlation between ACTH and corticosterone levels results demonstrated that corticosterone secretion following cold exposure is independent of ACTH (Y = 0.03x + 52.04, *r* = 0.07, *p* > 0.05).

## Discussion

The correlation between cortisol levels and blood collection site in the present study suggests there were differences in cortisol levels at the various collection sites tested. Therefore, we assessed the relationship between cortisol levels in blood collected from each site and cause of death and found that cortisol levels in cases of hypothermia were three times higher than those in other causes of death. No significant correlations were observed between cortisol levels and causes of death other than hypothermia. There was no correlation between ACTH concentration and cortisol levels in hypothermia, suggesting that cortisol can be produced by the adrenal gland during cold stress without stimulation by ACTH. Such ACTH-independent production of cortisol might be protective during prolonged (but not acute) periods of cold stress, as cold exposure promotes glucose production [55]. Importantly, micromorphologic changes in hormone expression in the adrenal cortex appear to be important for cold-induced cortisol secretion.

Cortisol is produced primarily in the zona fasciculata of the adrenal gland. Cell counts and nuclear and cytoplasm staining by technicians blinded to cause of death showed that during hypothermia, cortisol staining was primarily localized in the nucleus rather than the cytoplasm. Furthermore, nuclear stating of cortisol was significantly greater in cases of hypothermia than cases involving other causes of death, whereas no significant difference between groups was noted in terms of cytoplasmic staining. These findings support studies showing that glucocorticoid receptors are inactive in the cytoplasm, as they are complexed with other proteins [56]. When glucocorticoids bind, they become active dimers, move into the nucleus, and promote transcription. Here, we found high levels of cortisol staining in the nucleus during cold exposure. Considered together, these observations suggest that cortisol is secreted in large quantities in response to the stress of cold exposure and that re-uptake might also occur [57-59].

In this study, we used a novel co-culture system to assess ACTH and corticosterone secretion secondary to cold stimulation. We demonstrated that ACTH and corticosterone secretion levels and patterns differed and were not correlated. Mono-culture of ACTH- and corticosterone-secreting cells under ACTH-free conditions at 4°C resulted in a sudden peak in ACTH at 10 min that decreased after 30 min. This can be explained by the half-life of mouse ACTH [60]. However, in an ACTH-free environment, the increase in corticosterone was lower than that seen under co-culture conditions, and there was no correlation between corticosterone and ACTH levels. These results suggest that cold exposure leads to independent increased secretion of cortisol.

There are some limitations to this study. The correlation between ACTH and corticosterone levels in mouse cell culture may differ from that observed in human autopsy examples. The half-life of hormones may also differ in the cell culture models and in humans. Furthermore, it is necessary to examine differences between human cortisol and mouse corticosterone and address problems associated with temperature settings in the cell culture model [61].

In conclusion, the present study showed that serum cortisol level can be used as a biomarker for cold exposure and that cortisol production in response to cold stress does not depend on ACTH-based activation. As immunostaining for cortisol revealed high expression levels in the nucleus after cold exposure, it is possible that cortisol production following cold exposure is independent of ACTH stimulation.

## Supporting information

**Fig S1. Correlation with cortisol level.** Relationship between cortisol level in blood collected at different sites and sex (a), age (b), survival period (c), and postmortem period (d).

**Fig S2. Relationship between cortisol level and blood collection site.** Left and right cardiac blood (a), left cardiac blood-iliac vein blood (b), and right cardiac blood-iliac vein blood (c).

